# Growth under high light and elevated temperature affects metabolic responses and accumulation of health-promoting metabolites in kale varieties

**DOI:** 10.1101/816405

**Authors:** Sara Alegre, Jesús Pascual, Andrea Trotta, Peter J. Gollan, Wei Yang, Baoru Yang, Eva-Mari Aro, Meike Burow, Saijaliisa Kangasjärvi

## Abstract

Plants are highly sensitive to changes in the light environment and respond to alternating light conditions by coordinated adjustments in foliar gene expression and metabolism. Here we assessed how long-term growth under high irradiance and elevated temperature, a scenario increasingly associated with the climate change, affects foliar chemical composition of Brassicaceous plants. Transcript profiling of Arabidopsis suggested up-regulation of phenylpropanoid metabolism and down-regulation of processes related to biotic stress resistance and indole glucosinolates (GSL). These observations prompted metabolite profiling of purple (Black Magic) and pale green (Half Tall) varieties of kale, an economically important crop species. Long-term acclimation to high light and elevated temperature resulted in reduced levels of 4-methoxy-indol-3-yl-methyl GSL in both kale varieties. The total levels of aliphatic GSLs increased under these conditions, although the profiles of individual GSL structures showed cultivar-dependent differences. Black Magic became rich in 4-methylsulfinylbutyl GSL and 2-phenylethyl GSL, which have health-promoting effects in human diet. Additionally, the purple pigmentation of Black Magic became intensified due to increased accumulation anthocyanins, especially derivatives of cyanidin. These findings demonstrate that the potentially stressful combination of high light and elevated temperature can have beneficial effects on the accumulation of health-promoting metabolites in leafy vegetables.

## Introduction

Light is an important external factor that drives photosynthesis, metabolism and growth in plants. To cope with varying light conditions, plants undergo coordinated acclimation responses, which can occur across molecular, cellular and whole plant levels and range from fast photosynthetic rearrangements to durable adjustments in metabolite composition, morphology, flowering time and seed production (Aro *et al*., 1993; Foyer, 2018; Pascual *et al*., 2017). Studies on model plants, notably *Arabidopsis thaliana* (hereafter Arabidopsis), have elucidated the dynamic nature of gene expression occurring upon short-term fluctuations in light conditions (Spetea, Rintamäki & Schoefs 2014; Gollan, Tikkanen & Aro 2015; Crisp *et al*. 2017). In nature, high light is commonly accompanied by elevated temperature, and episodes of bright and hot conditions may become more frequent due to climate change. A typical response to the potentially stressful combination of light and heat is accumulation of carotenoids and phenolic pigments, which can protect foliar tissues against light-induced damage (Chalker-Scott 1999; Zeng, Chow, Su, Peng & Peng 2010). Long-term metabolic responses to high light and elevated temperature, beyond regulation of photosynthesis and chloroplast metabolism, have remained less well understood.

Sulphur metabolism is tightly linked with light-driven redox chemistry in chloroplasts and yields a number of precursors, metabolic intermediates and specialised compounds, which are vital in mediating defensive responses upon environmental challenges (reviewed by Chan *et al*., 2019). In cruciferous plants, sulphur metabolism sustains the biosynthesis of glucosinolates (GSLs), which are sulfur- and nitrogen-containing specialised metabolites whose breakdown products cause the characteristic pungent taste of Brassica crops (Bell, Oloyede, Lignou, Wagstaff & Methven 2018). Studies on Arabidopsis, kale (*Brassica oleracea* convar *acephala*), broccoli (*B. oleracea var. italica*) and cabbage (*B. oleracea var. capitata*) have elucidated the commercial and ecological impacts of GSLs in human and animal nutrition as well as in plant-environment interactions, such as those with pathogens and herbivores (Wittstock & Burow 2010; Sharma, Singh & Mikawlrawng 2014; Traka 2016; Francisco *et al*. 2017). In the human diet, consumption of GSL-rich cruciferous crops has been associated with a reduced risk of cancer (Gupta, Wright, Kim & Srivastava 2015; Megna, Carney, Nukaya, Geiger & Kennedy 2016; Katz, Nisani & Chamovitz 2018) and chronic inflammation diseases (Sun *et al*. 2015; Yamagishi & Matsui 2016), but certain GSL species might also have detrimental effects in animal nutrition (Felker, Bunch & Leung 2016). Kale breeding has yielded a multitude of varieties that differ with respect to shape, coloration and the content of specialized metabolites, which can increase the nutritional and market value of the leafy vegetables (Bell *et al*. 2018).

Studies on Arabidopsis have elucidated the biosynthesis, modification, degradation and transport of certain GSLs (Halkier & Gershenzon 2006; Sønderby, Geu-Flores & Halkier 2010; Jensen, Halkier & Burow 2014) and uncovered mechanisms behind transcriptional and post-translational regulation of these processes (Celenza *et al*. 2005; Gigolashvili *et al*. 2007; Frerigmann *et al*. 2016; Rahikainen *et al*. 2017). The GSL core structure consists of a glucose moiety bound to a sulfonated aldoxime and a variable amino acid-derived side chain. The structural diversity of GSLs species stems from modifications that may take place in both the side group and the core structure (Sønderby *et al*. 2010; Jeschke & Burow 2018). Recently, formation of a methoxylated tryptophan-derived indole GSL, 4-methoxy-indol-3-yl-methyl GSL (4MO-I3M GSL), was functionally connected with *S-*Adenosyl-L-Homocysteine Hydrolase (SAHH), which is the key enzyme of the activated methyl cycle, essential for all trans-methylation reactions in all living cells (Rahikainen, Alegre, Trotta, Pascual & Kangasjärvi 2018). In Arabidopsis, accumulation of SAHH in distinct oligomeric complexes correlated with increased abundance of 4MO-I3M, but the potential role of SAHH in defence and its link to growth light conditions remain obscure (Rahikainen *et al*. 2017).

Here we explored how growth under high light and elevated temperature affects metabolic adjustments in kale, an economically relevant brassicaceous crop species. Analysis of transcriptomic datasets available for Arabidopsis suggested up-regulation of processes related to phenolic compounds, while processes related to biotic stress resistance and indole GSL became down-regulated. These observations prompted analysis of amino acids, anthocyanins and GSL contents in kales, which revealed both stress-induced and cultivar-dependent adjustments in purple (Black Magic, BM) and pale green (Half Tall, HT) varieties of this leafy vegetable. Both kale varieties responded to long-term growth under high light and elevated temperature by reducing the contents of methionine, the methyl donor *S*-adenosyl methionine (SAM) and the methoxylated indole GSL 4MO-I3M. In contrast, the total contents of methionine-derived aliphatic GSLs increased in the high-light-grown kales, with distinct cultivar-dependent GSL-profiles. When grown under the warm high light conditions, Black Magic became particularly rich in specific anthocyanins and health-promoting aliphatic GSLs. Collectively, translation of the basic knowledge from Arabidopsis to kales highlighted the effect of growth light on the foliar chemical composition and nutritional properties of leafy vegetables.

## Material and Methods

### Plant material

*Arabidopsis thaliana* (L.) Heynh. ecotype Columbia-0 was grown in 50% relative humidity and 8/16-hour photoperiod under growth light (GL; 130 µmol photons m^−2^ sec^−1^ and 22°C) for 2 weeks and thereafter shifted for acclimation under high light (HL; 800 µmol photons m^−2^ sec^−1^ and 28°C) for 2 weeks. Control plants were grown under GL conditions for 4 weeks. *Brassica oleracea* convar. *acephala,* Half Tall and Black Magic, were grown in 50% relative humidity and 12/12-hour photoperiod. Plants were grown under growth light (130 µmol photons m^−2^ sec^−1^ and 22°C) or in high light (800 µmol photons m^−2^ sec^−1^ and 26°C). Plants were germinated in GL and transferred to one of the above-mentioned conditions two days after germination. Experiments with kales were carried out with 19 days old plants. For each kale variety and condition leaves of 16 plants were collected. Each biological replicate consisted of the longitudinal halves of leaves from two plants, resulting in eight biological replicates.

### Analysis of gene expression

Rosettes of Arabidopsis grown under GL or acclimated for two weeks in HL were collected four hours after the onset of light period and analysed by Agilent Arabidopsis (V4) Gene Expression Microarrys, 4×44K (Design ID 021169) as described by Konert *et al*., 2015.

### Comparison of gene expression profiles

Genes with absolute expression fold change (FC) >2 and p-values <0.05 in the long-term Arabidopsis HL dataset presented in Supplementary Table S1 were compared with publicly-available datasets obtained from short-term shifts to HL. The short-term HL treatments included in the analysis are listed in Supplementary Table S2. They were selected by comparing the 400 most significantly differentially expressed genes in the long-term HL dataset (Supplementary Table S1) with the Affymetrix Arabidopsis ATH1 Genome Array database, querying against experiments containing the keyword “high light” in the Genevestigator (RRID:SCR_002358) database (Hruz *et al*. 2008).

The RAW data of the selected short-term HL experiments (summarized in supplementary Table S2) were downloaded from Gene Expression Omnibus (RRID:SCR_005012; https://www.ncbi.nlm.nih.gov/geo/) and ArrayExpress (RRID:SCR_002964; https://www.ebi.ac.uk/arrayexpress/) and pre-processed independently in Bioconductor (RRID:SCR_006442; http://www.bioconductor.org/), comprising normalization by Robust Multi-array Average. Differential gene expression was analyzed by *limma* package (v3.36.5; RRID:SCR_010943) (Smyth 2004) using the Benjamini-Hochberg false discovery rate for adjusting p-values for multiple hypothesis testing. Absolute FC>2 and p-values <0.05 were considered differential. The transcript profiles were hierarchically clustered with R package *pheatmap* (v1.0.12) (Kolde 2019) using Ward’s method and Euclidean distance.

Venn diagram was generated with VennDiagram R package (v1.6.20; RRID:SCR_002414) (Chen & Boutros 2011). All analyses were performed in R (v3.5.1) (RCore Team, 2018) run in RStudio (v1.1.456; RRID:SCR_000432) (RStudio Team, 2018). Gene Ontology enrichment analysis were performed and GO trees were generated with ShinyGO v0.60 (Ge & Jung 2018).

### Photosynthesis measurements

Photosynthetic activity was estimated by analysis of chlorophyll *a* fluorescence and P700 oxidation using a DUAL-PAM 100 measuring system (Waltz, Germany). The apparent electron transport rates of PSII and PSI were derived from the calculated quantum yields, according to the formula ETR = yield × PAR × 0.84 × 0.5, where 0.84 is the average radiation absorbed by the leaf and 0.5 is the fraction of photons distributed to each photosystem. Non-photochemical quenching (NPQ) was calculated by NPQ = (Fm–Fm′)/Fm′. ETR(I), ETR(II) and NPQ were assessed increasing light intensities of 50, 125, 500 and 1000 µmol photons m^−2^ s^−1^ in leaves after 30 min in darkness.

### Spectrophotometric measurement of total leaf pigments

Spectrophotometric quantification of kale leaf pigments was performed as in Sims and Gamon, 2002. Carotenoids were extracted from 100 mg of frozen leaf powder with 400 µl acetone/tris buffer solution (80:20 v/v, pH 7.8). Anthocyanin contents were analyzed using 100 mg of powder extracted in acidified methanol (0.1% HCl, v/v).

### Mass spectrometric analysis of anthocyanins

Anthocyanins were extracted from 500 mg of frozen leaf powder in a total volume of 45 ml of acidified methanol (0.1% HCl, v/v). After centrifugation, the supernatant was collected, the organic solvent was removed using a vacuum rotary evaporator at 40°C. The samples were dissolved in 1 ml of acidified methanol (1% HCl, v/v). Qualitative analysis of anthocyanins was performed using a Waters Acquity ultrahigh-performance liquid chromatography (UPLC) system (Waters Corp., Milford, MA, USA) combined with Waters Quattro Premier Tandem Quadrupole mass spectrometer (Waters Corp., Milford, MA, USA) equipped with an electrospray ionization (ESI) source. A Phenomenex Aeris peptide XB-C18 (3.6 μm, 150 × 4.60 mm) column combined with a Phenomenex Security Guard Cartridge Kit (Torrance, CA, USA) was used and maintained at 35°C during the analysis. The analyses were carried out by a gradient elution with formic acid/water (5:95, v/v) as solvent A and acetonitrile as solvent B at a flow rate of 1 mL/min. The gradient program of solvent B in A (v/v) was 0–1 min with 4–6% B, 1–2 min with 6–8% B, 2–6 min with 8–9% B, 6–10 min with 9–10% B, 10–15 min with 10–20% B, 15–20 min with 20–25% B, 20–25 min with 25–80% B, 25–30 min with 80–4% B, and 30–35 min with 4% B. The injection volume was 10 μL. A split joint was applied and directed a flow of 0.4 mL/min into the mass spectrometer after the UV detector. The ESI-tandem mass spectrometry (MS/MS) was operated according to the previous method by Yang *et al*., 2018.

Quantitative analysis of anthocyanins was carried out using a Shimadzu Nexera UHPLC system (Shimadzu Corporation, Kyoto, Japan), which consisted of a CBM-20A central unit, a SIL-30AC auto sampler, two LC-30AD pumps, a CTO-20AC column oven, and a SPD-M20A diode array detector. The chromatographic conditions were the same as described above in qualitative analysis. An external standard of cyanidin 3-*O*-glucoside was used for quantitative analysis, and all the anthocyanins were quantified as equivalents of cyanidin 3-*O*-glucoside using the calibration curve constructed with this reference compound. The total content of anthocyanins was calculated as the sum of the peaks, which represented a minimum of 1% of the total peak area in the chromatogram.

### Glucosinolate analysis

Frozen fresh material was homogenized in a bead mill (two 3 mm chrome balls, 2 × 30 s at 30 Hz) and ~150 mg subsequently extracted with 1 mL ice-cold 85% (v/v) methanol containing 20 nmol p-hydroxybenzyl glucosinolate (pOHb; PhytoLab, cat. No. 89793) as internal standard. After centrifugation (10 min, 13.000 × *g*, 4°C), GSLs were extracted from 150 µl of the supernatant as desulfo-GSLs as described before (see Alternate Protocol 2, Crocoll *et al*., 2016). LC-MS/MS analysis was carried out on an Advance UHPLC system (Bruker, Bremen, Germany) equipped with C18 column (Kinetex 1.7 u XB-C18, 10 cm × 2.1 mm, 1.7 µm particle size, Phenomenex, Torrance, CA, USA) coupled to an EVOQ Elite TripleQuad mass spectrometer (Bruker, Bremen, Germany) equipped with an electrospray ionisation source (ESI). The injection volume was 1 µL. Separation was achieved with a gradient of water/0.05% (v/v) formic acid (solvent A) - acetonitrile (solvent B) at a flow rate of 0.4 mL/min at 40°C (formic acid, Sigma-Aldrich, cat. no. F0507; acetonitrile). The elution profile was 0-0.5 min with 2% B; 0.5-1.2 min with 2-30% B; 1.2-2.0 min with 30-100% B; 2.0-2.5 min with 100% B; 2.5-2.6 min with 100-2% B; 2.6-4.0 min with 2% B. The ion spray voltage was maintained at +3500 V. Cone temperature was set to 300°C and cone gas to 20 psi. Heated probe temperature was set to 400°C and probe gas flow set to 40 psi. Nebulising gas was set to 60 psi and collision gas to 1.6 mTorr. Desulfo-GLSs were monitored based by Multiple Reaction Monitoring (MRM) with appropriate analyte parent ion to product ion transitions as previously described (Crocoll *et al*. 2016). Quantification of the individual GLSs was based on response factors relative to the internal standard pOHB calculated from standard curves in control extracts.

### Amino acid analysis

A 20-µl aliquot of the supernatant collected as described for glucosinolate analysis was diluted 1:10 (v/v) mixed with a stock solution containing 10μg/mL ^13^C-, ^15^N-labelled amino acids (Algal amino acids ^13^C, ^15^N, Isotec, Miamisburg, US). The resulting samples were filtered (Durapore®0.22μm PVDF filters, Merck Millipore, Tullagreen, Ireland) and used directly for LC-MS/MS analysis. Chromatography was performed on an Advance UHPLC system (Bruker, Bremen, Germany) with a Zorbax Eclipse XDB-C18 column (100×3.0 mm, 1.8 μm, Agilent Technologies, Germany). Formic acid (0.05% (v/v) in water and acetonitrile (supplied with 0.05% (v/v) formic acid) were employed as mobile phases A and B, respectively. The elution profile was: 0–1.2 min with 3% B; 1.2–4.3 min with 3–65% B; 4.3–4.4 min with 65–100% B; 4.4–4.9 min with 100% B, 4.9–5.0 min with 100-3% B and 5.0–6.0 min with 3% B. Mobile phase flow rate was 500 μl/min and column temperature was maintained at 40°C. LC was coupled to an EVOQ Elite Triple Quad mass spectrometer (Bruker, Bremen, Germany) equipped with electrospray ionisation. The ion spray voltage was maintained at 3000 V or −4000 V in positive or negative ionisation mode, respectively. Cone temperature was set to 300°C and cone gas flow to 20psi. Heated probe temperature was set to 400°C and probe gas flow set to 50 psi. Nebulising gas was set to 60 psi and collision gas to 1.6m Torr. Nitrogen was used as both cone gas and nebulizing gas and argon as collision gas. MRMs for the ^13^C, ^15^N-labelled amino acids were chosen as previously described (Docimo *et al*. 2012). Response factors for quantification of amino acids, SAM and cystathione had been calculated previously based on dilution series of the respective analytes (Petersen, Crocoll & Halkier 2019).

### Statistical analyses of metabolite data

All the statistical analysis were performed in R environment v 3.5.1 (RRID:SCR_000432). Numerical data obtained from analysis of amino acids, GSLs and total pigments were subjected to statistical analysis using one-way ANOVA with statistical significance at the level of P < 0.05, followed by Tukey’s comparison in case distributions followed normality and homoscedasticity, whereas Kruskal-Wallis test was applied in the rest of the cases.

Black Magic anthocyanins content was analyzed with Student’s T-test and significance level of P< 0.05 is denoted by an asterisk.

## Results

### Transcript profiling of Arabidopsis leaves after long-term acclimation to high light and elevated temperature

To gain insights into how growth under high light and moderately elevated temperature might affect metabolic processes in brassicaceous plants, we took advantage of the genetic resources available for Arabidopsis. First we performed microarray analysis of Arabidopsis grown under moderate growth light and moderated temperature (GL; 130 µmol photons m^−2^s^−1^/22°C) or acclimated to high light and elevated temperature (HL+ET; 800 µmol photons m^−2^s^−1^/28°C) (Figure 1a). Arabidopsis responded to a two-week growth period under HL+ET by visually observable accumulation of purple pigments (Figure 1a). To determine the effects of long-term HL+ET acclimation on gene expression, genes with >2 FC (p-value <0.05) differential expression in plants acclimated to HL+ET were identified, in comparison to plants grown in GL (Figure 1b; Table S1). Gene Ontology (GO) enrichment among differentially expressed genes revealed that the main processes up-regulated in response to long-term HL+ET acclimation included anthocyanin biosynthesis and metabolism, flavonoid metabolism and abiotic stress responses (Figure 1b). Genes related to photosynthesis and light harvesting, in contrast, were among the most down-regulated. Notably, GO categories related to defense responses, salicylic acid (SA) signaling and indole GSL metabolism were also over-represented among the down-regulated genes, when compared to plants grown in GL (Figure 1b; Table S1).

**Figure 1.**
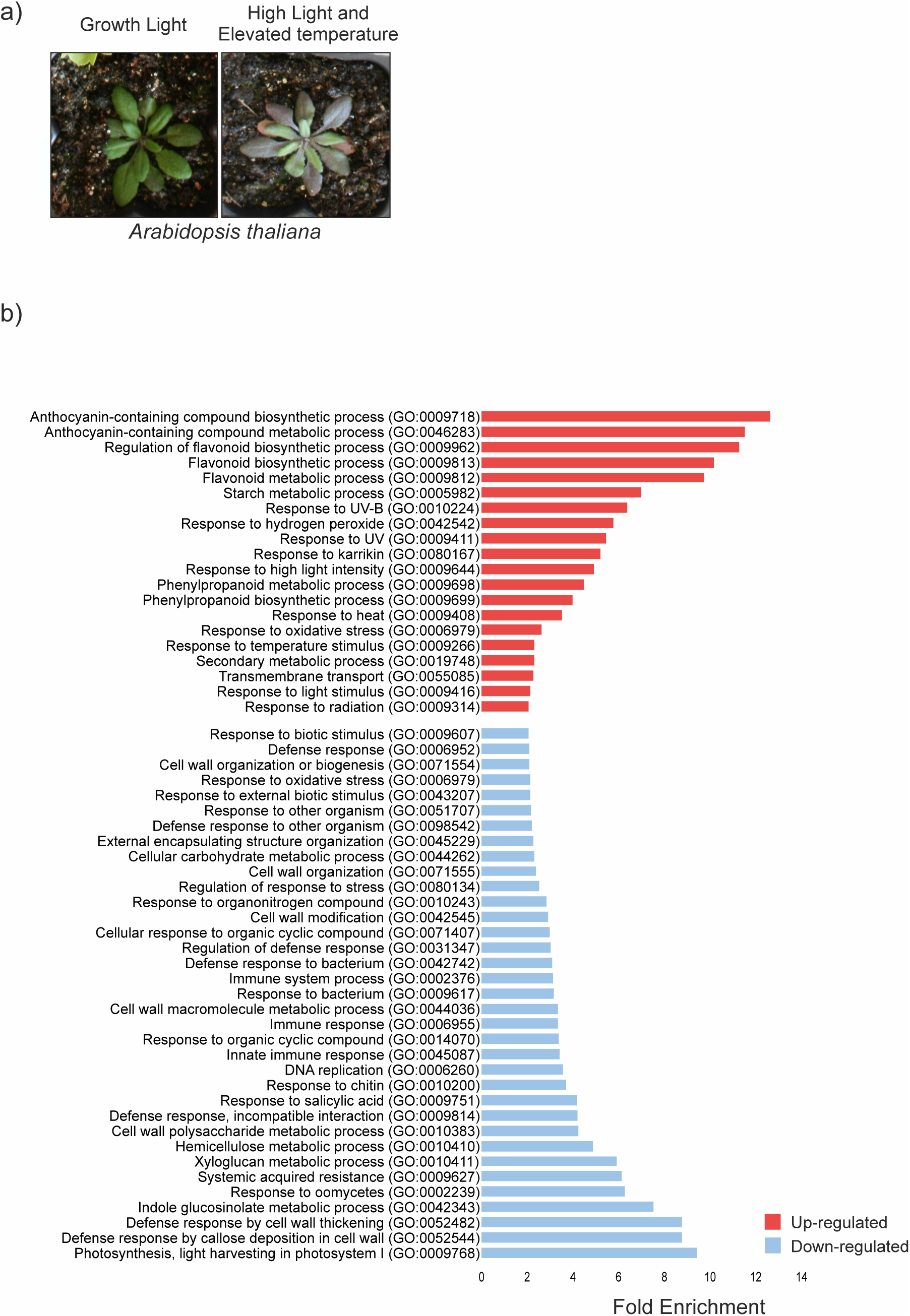
Phenotypic characteristics and gene ontology (GO) categories enriched in the transcript profile of Arabidopsis wild type acclimated to long-term high light and elevated temperature. Two-week-old *Arabidopsis thaliana* wild type was shifted from growth light (130 µmol photons m^−2^ sec^−1^ and 22°C) to high light and elevated temperature (HL+ET; 800 µmol photons m^−2^ sec^−1^ and 28°C) for 2 weeks. A) Phenotypic characteristics of *Arabidopsis thaliana* after long-term acclimation to high light and elevated temperature. B) Gene Ontology (GO) categories enriched in the transcript profile of high light and elevated temperature-acclimated *Arabidopsis thaliana* wild type plants, when compared to plants grown under growth light and moderated temperature.

Next we utilized publicly available Arabidopsis transcriptomic datasets to determine how the effects of long-term HL+ET exposure differ from short-term HL treatments. Transcripts with >2 FC (p-value <0.05) differential expression in the long-term HL+ET-acclimated plants (Supplementary Table S1) were selected and their abundance was assessed in the publicly available datasets obtained from plants exposed to various short-term HL treatments (summarized in Supplementary Table S2), ranging from 30 minutes to 6 hours as detailed in Supplementary Table S3.

Hierarchical clustering analysis grouped the short term treatments into two clusters, early time points (30 min, 1 h and 2 h) and later ones (3h, 4h and 6 h) (Figure 2). The analysis also revealed temporal HL-induced transcriptional responses, which formed seven main clusters (Figure 2; Table S3). Cluster 1 contained transcripts whose abundance became reduced already at early time points of HL illumination. This early-responding cluster was enriched in GO terms related to biotic stress and cell wall metabolism (Table S4; Figure S1a). Clusters 2 and 3 comprised of transcripts that showed reduced abundance almost exclusively in the long-term HL dataset (Tables S5 and S6; Figure S1b,c). Clusters 4 and 6 in turn included transcripts with increased abundance upon long-term HL acclimation (Table S8; Figure S1e). Clusters 5 and 7 comprised transcripts whose abundance was increased in the long-term high light dataset but varied between the short-term HL (Tables S7 and S9; Figure S1d,f).

**Figure 2.**
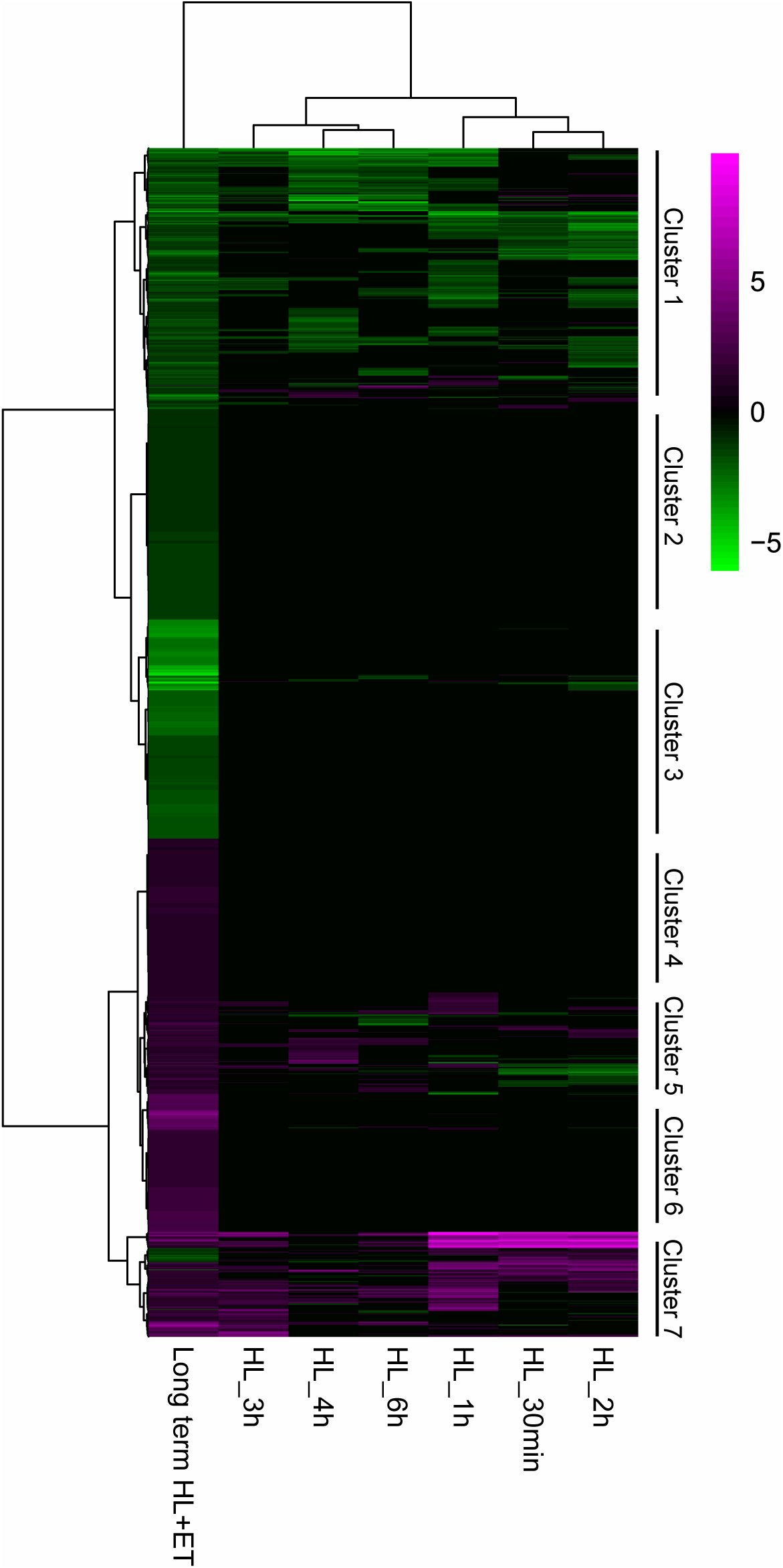
Hierarchical clustering of gene expression profiles from long-term high light- and-elevated-temperature-acclimated plants and plants exposed to short-term high light shifts. Purple denotes upregulation and green denotes downregulation of transcript abundance. The long-term high light acclimation dataset was obtained in this study (Table S1), while the others were downloaded from AtGenExpress and Gene Expression Omnibus (see Supplementary Tables S2 and S3 for full description of the datasets).

Comparison of the sets of genes differentially expressed in the long-term and representative short-term HL treatments identified 49 genes differentially expressed in response to every HL treatment (marked in blue in Table S3) and 1469 genes, which were differentially expressed exclusively upon acclimation to long-term HL+ET, but not in any of the short-term light shifts (Figure 3; Table S3). Within this group, GO enrichment analysis of transcripts with increased abundance specifically in long-term HL+ET revealed over-representation of categories related to transcriptional regulation, membrane transport and regulation of biosynthetic processes and biosynthesis of flavonoids (Table S10; Figure S2a). Among transcripts with reduced abundance specific to long-term HL+ET acclimation, GO categories related to chlorophyll biosynthesis, photosynthetic light-harvesting, DNA integrity and biotic stress responses were significantly over-represented (Table S11; Figure S2b).

**Figure 3.**
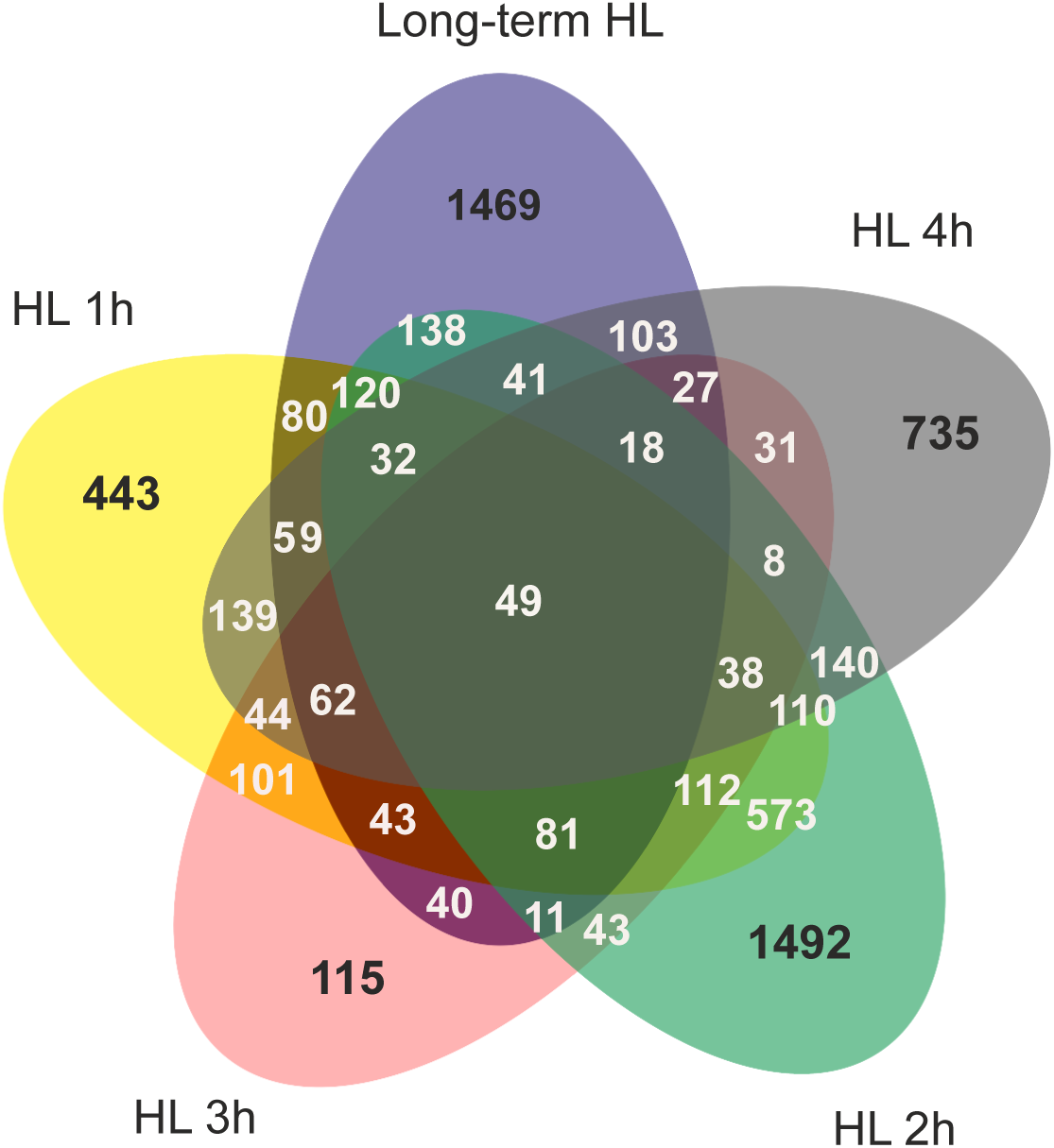
Venn diagram depicting overlaps between the sets of differentially accumulated transcripts in long-term high light and elevated temperature and high light shift experiments according to the hierarchical clustering presented in. Figure 2.

### Light intensity-dependent phenotypic characteristics in differentially pigmented kale varieties

The enrichment of genes related to anthocyanins and GSL in the Arabidopsis long-term HL+ET transcriptome (Figure 1) prompted us to assess the effect of growth conditions on the contents of these nutritionally important compounds in kales, which are commercially important leafy vegetables. Two varieties of *Brassica oleracea* convar. *acephala*, Half Tall and Black Magic with differential pigmentation patterns, were selected for the analysis. Growing the kales under 800 µmol photons m^−2^ sec^−1^ at 28°C triggered typical high-light-induced morphological responses, such as shorter petioles, reduced height and thicker leaves (Figure 4). In Black Magic, the light avoidance response was additionally evident as vertical disposition of the leaves, while Half Tall showed a twisted leaf morphology when grown under HL (Figure 4).

**Figure 4.**
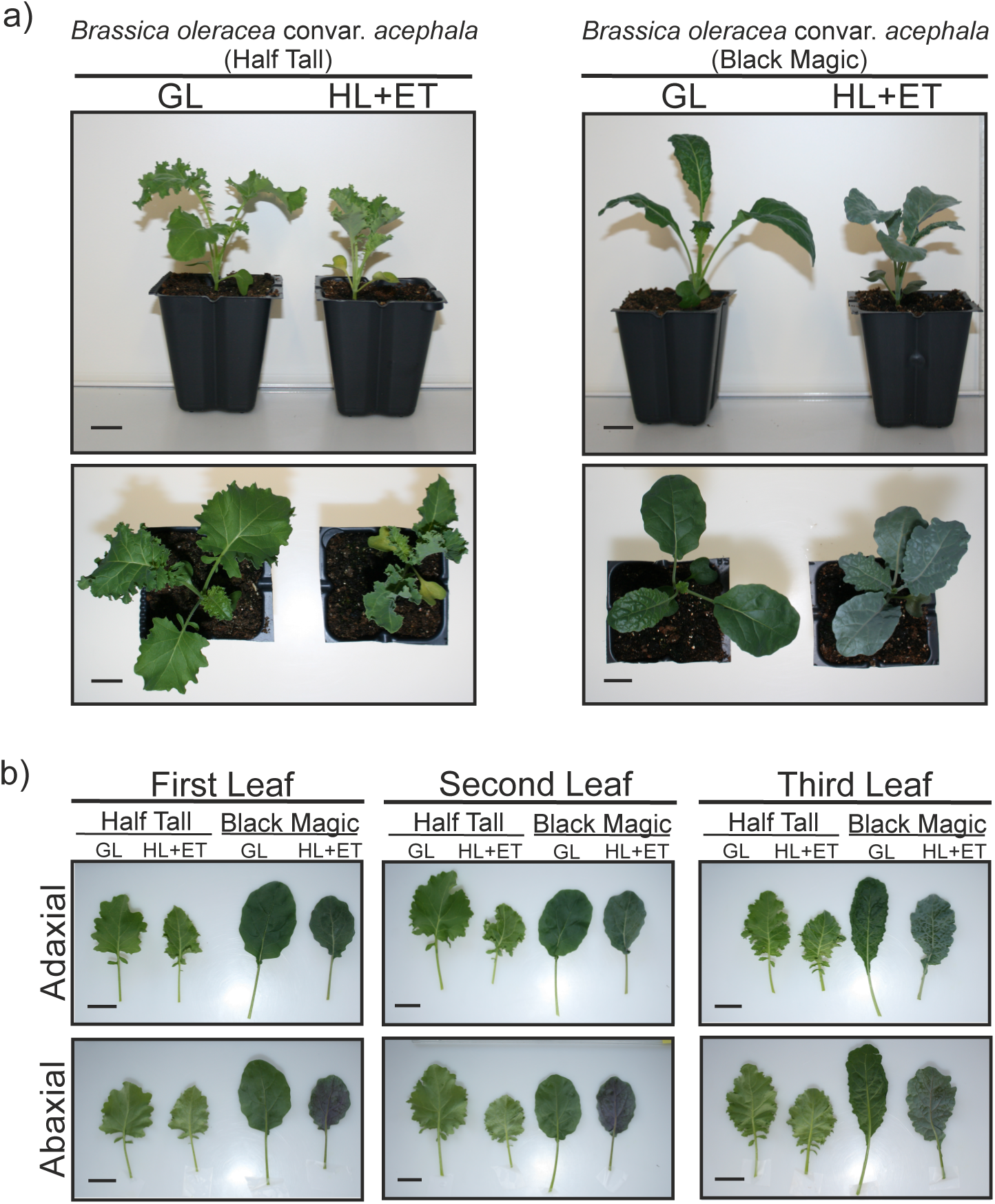
Visual characteristics of the kale varieties *Brassica oleracea* convar. *Acephala* Half Tall and Black Magic. A) Morphological characteristics of Half Tall (HT) and Black Magic (BM) after 3 weeks of growth under 130 µmol photons m^−2^s^−1^ at 22°C (GL) or 800 µmol photons m^−2^s^−1^ at 26°C (HL+ ET). Scale bar corresponds to 2 cm. B) Representative photographs depicting light- and temperature-dependent adjustments in leaf morphology and pigmentation as visualized from adaxial and abaxial surface of the first, second and third leaves. Scale bar corresponds to 2 cm.

The effect of HL+ET acclimation on the photosynthetic capacity of the two kale varieties was assessed by comparing the performance of the photosynthetic light reactions between Black Magic and Half Tall grown in either GL or HL+ET, using a DUAL-PAM-100. The rates of photosynthetic electron transport (ETR) through PSII and PSI were higher in Black Magic leaves from both GL and HL+ET, in comparison to Half Tall leaves, at actinic irradiances above 125 µmol photons m^−2^ s^−1^ (Figure 5a,b). Furthermore, ETR(II) and ETR(I) in HL+ET-grown plants were higher than their GL-grown counterparts at 500 and 1000 µmol photons m^−2^ s^−1^ actinic light. Leaves from HL+ET-grown plants demonstrated significantly lower NPQ at 50 µmol photons m^−2^ s^−1^ than GL-grown plants, while NPQ in HL+ET-grown Black Magic leaves was also significantly lower than all other samples at 125 µmol photons m^−2^ s^−1^. At 500 and 1000 µmol photons m^−2^ s^−1^, Half Tall leaves had higher NPQ than Black Magic leaves, irrespective of growth conditions, while NPQ did not differ significantly between GL- and HL+ET-grown plants of each variety at these irradiances (Figure 5c).

**Figure 5.**
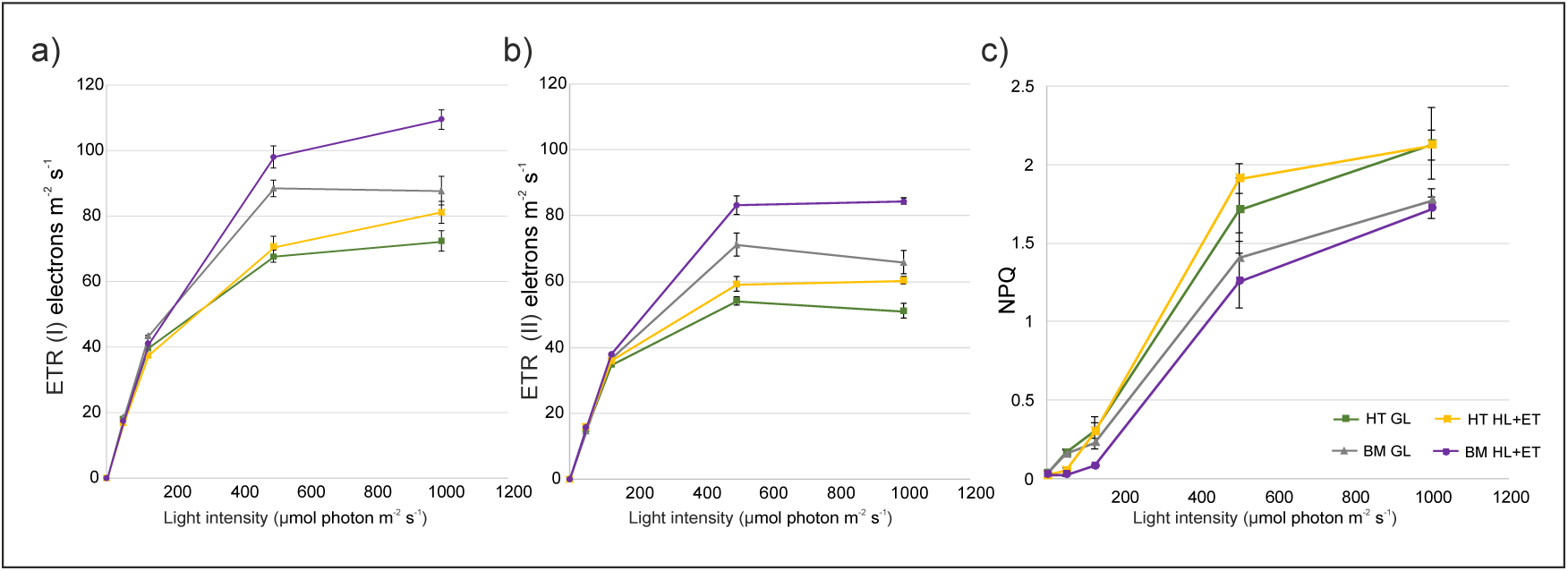
Photosynthetic performance of differentially light -acclimated kales. Half Tall (HT) and Black Magic (BM) were grown for 3 weeks under 130 µmol photons m^−2^s^−1^ at 22°C (GL) or 800 µmol photons m^−2^s^−1^ at 26°C (HL+ET) and the photosynthetic parameters were measured with Dual-Pam. A) **ETR (I),** Electron transport rate. B) **ETR (II),** Electron transport rate. C) **NPQ,** Non-photochemical quenching.

The purple pigmentation of Black Magic became intensified upon growth under HL+ET and accumulation of protective pigments was evident on both adaxial and abaxial sides of the leaves (Figure 4b). Spectrophotometric analysis further indicated elevated amounts of both anthocyanins and carotenoids in HL+ET-acclimated Black Magic leaves (Figure 6). Half Tall, in contrast, was devoid of this common protective response and did not undergo light-induced accumulation of these pigments (Figure 6).

**Figure 6.**
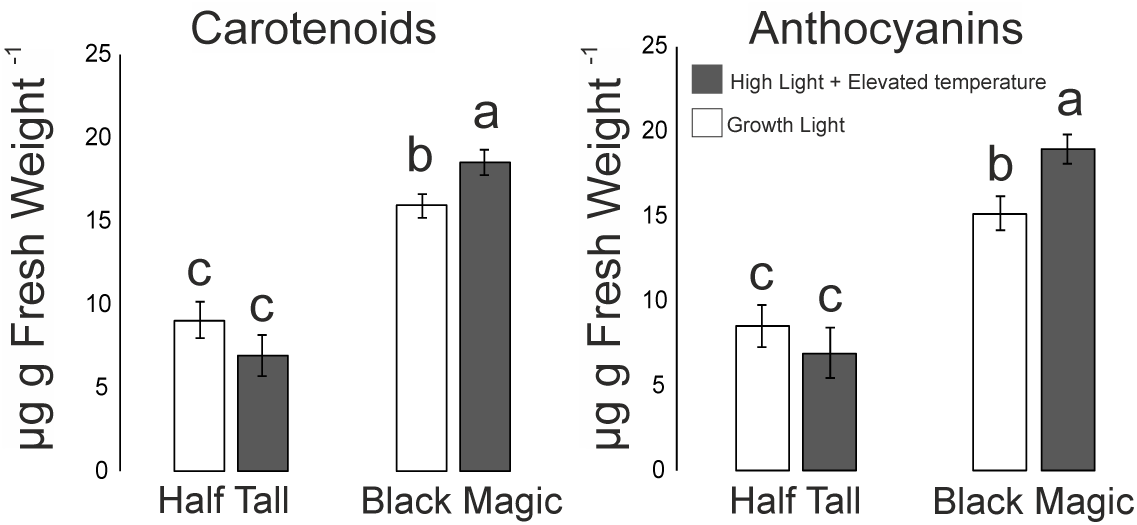
Total content of carotenoids and anthocyanins in differentially light-acclimated Half Tall and Black Magic leaves. Half Tall (HT) and Black Magic (BM) were grown for 3 weeks under 130 µmol photons m^−2^s^−1^ at 22°C (GL) or 800 µmol photons m^−2^s^−1^ at 26°C (HL+ET) and the pigments were quantified by spectrophotometry. Error bars show the SE and different letters indicate statistically significant differences (p< 0.05), n=4.

For more detailed assessment of anthocyanin contents, kale leaf extracts were analyzed by UPLC-ESI-MS/MS and the detected anthocyanins were qualitatively described based on mass spectra, UV spectra and comparison to previously published literature (Olsen, Aaby & Borge 2010; Olsen, Grimmer, Aaby, Saha & Borge 2012) (Table 1). In Black Magic, ten different compounds were detected (Figure S3). The anthocyanins existed in different acylated forms with sinapic acid, ferulic acid, caffeic acid and *p*-coumaric acid as the predominant acyl donors (Table 1). Compounds 9 and 10, identified as cyanidin-3-sinapoyl-feruloyl-diglucoside-5-glucoside and cyanidin-3-disinapoyl-diglucoside-5-glucoside, respectively, were the most abundant anthocyanins detected in Black Magic (Table 1). In Half Tall, only trace amounts of anthocyanins were detected and individual compounds could therefore not be reliably identified. Quantification of total anthocyanins revealed that the generally higher levels in Black Magic significantly increased upon acclimation to HL+ET (Figure 7).

**Figure 7.**
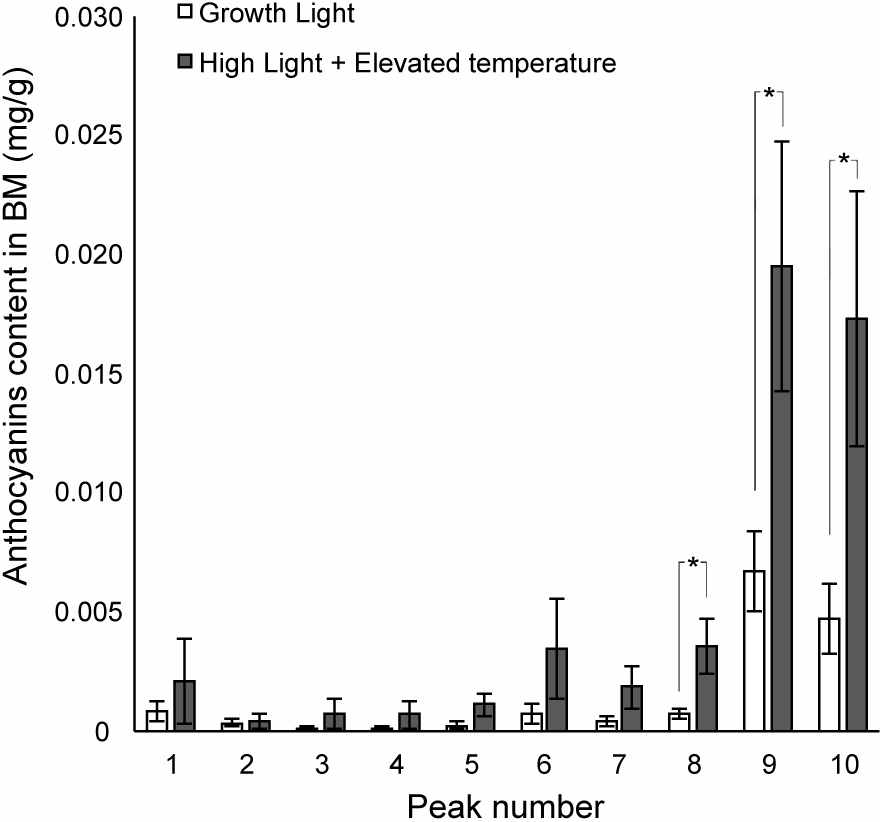
Anthocyanin profiles in differentially light acclimated Black Magic kales. Black Magic (BM) was grown for 3 weeks under 130 µmol photons m^−2^s^−1^ at 22°C (GL) or 800 µmol photons m^−2^s^−1^ at 26°C (HL+ET) and the pigments were quantified by tandem UPLC with mass spectrometry. On the X-axis, the numbers correspond to detected peaks, which are depicted in Figure S3. Error bars indicate SE and * indicate statistically significant differences (p< 0.05) between light condition for each different compound, n=4. Tentative identities of the compounds are listed in Table 1.

**Table 1.** Identification of anthocyanins from Black Magic leaf extracts.

| No<br>. | RT<sub>min</sub> | [M+<br>H]<sup>+</sup> | Daughter ions<br>(MS<sup>2</sup>) | Tentative ID |
| --- | --- | --- | --- | --- |
| 1 | 9.182 | 979 | 817, 449, 287 | cyanidin-3-sinapoyl-diglucoside-5-glucoside |
| 2 | 9.964 | 949 | 449, 287 | cyanidin-3-feruloyl-diglucoside-5-glucoside |
| 3 | 14.72<br>0 | 1288 | 1126, 617,<br>449, 287 | unknown |
| 4 | 14.95<br>2 | 1141 | 979, 653, 449,<br>287 | cyanidin-3-sinapoyl-caffeoyl-diglucoside-5-glucoside I |
| 5 | 15.88<br>1 | 1141 | 979, 653, 449,<br>287 | cyanidin-3-sinapoyl-caffeoyl-diglucoside-5-glucoside II |
| 6 | 16.16<br>2 | 949 | 449, 287 | cyanidin-3-caffeoyl-feruloyl-diglucoside |
| 7 | 16.81<br>6 | 1125 | 963, 449, 287 | cyanidin-3-sinapoyl-*p*-coumaroyl-diglucoside-5-glucoside |
| 8 | 16.93<br>0 | 1111 | 930, 287 | unknown |
| 9 | 17.09<br>0 | 1155 | 993, 449, 287 | cyanidin-3 –feruloyl-sinapoyl -diglucoside-5-glucoside |
| 10 | 17.35<br>9 | 1185 | 1023, 449, 287 | cyanidin-3-disinapoyl-diglucoside-5-glucoside |

### Light-induced changes in amino acids in kale leaves

Amino acid metabolism provides precursors for the biosynthesis of complex chemical structures, including some classes of specialized metabolites. Analysis of amino acids revealed an overall similarity between the amino acid profiles of Half Tall and Black Magic, with the exception of proline and cysteine, the contents of which differed between the varieties (Figure 8). Both Half Tall and Black Magic responded to HL+ET acclimation by diminished amounts of several amino acids. The levels of aspartic acid and some of its derivatives, including asparagine, lysine, threonine, isoleucine and methionine, as well as the contents of the metabolic intermediates L-cystathionine and SAM, became reduced in both kale varieties (Figure 8). Similarly, the contents of glutamate, arginine, alanine, serine, glycine and histidine declined under high light. In contrast, the contents of proline and leucine showed a trend towards increased levels in both varieties when the plants grew under HL+ET (Figure 8). The contents of phenylalanine, the precursor for the biosynthesis of phenolic pigments and aromatic GSLs did not vary between the treatments (Figure 8).

**Figure 8.**
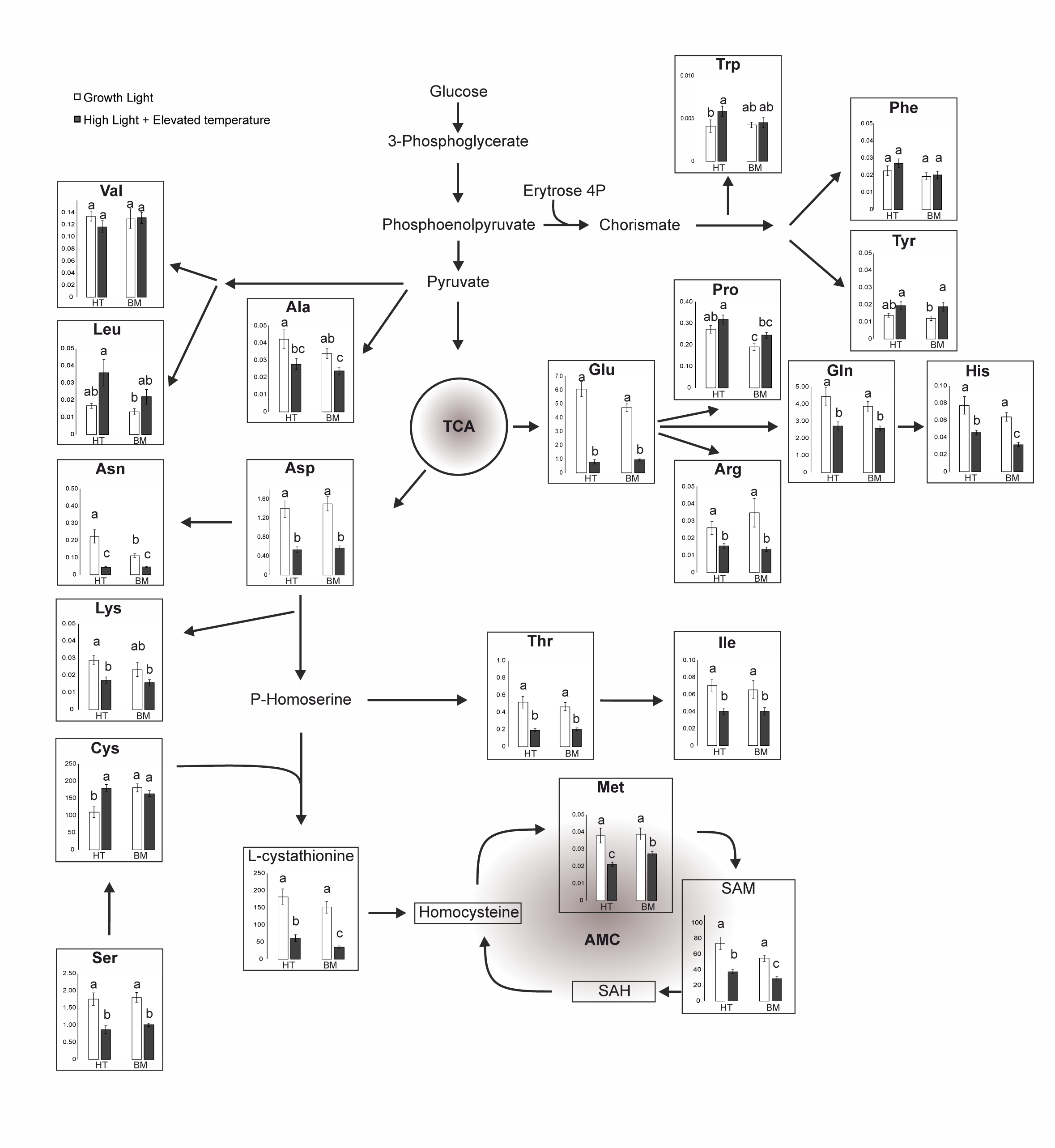
Contents of amino acids and SAM in differentially light-acclimated kales. Half Tall (HT) and Black Magic (BM) were grown under 130 µmol photons m^−2^s^−1^ at 22°C (GL) or 800 µmol photons m^−2^s^−1^ at 26°C (HL+ET) and the contents of amino acids and SAM were analysed LC-MS/MS. Quantitative values are expressed in nmol mg^−1^ FW. Error bars indicate SE and different letters indicate statistical significant differences (p< 0.05) n=8.

### SAHH complex formation in differentially light-acclimated kales

The HL+ET-induced reduction in the levels of methionine and SAM (Figure 8) suggested alterations in the activated methyl cycle, where SAHH is the key enzyme responsible for the maintenance of trans-methylation capacity (Rahikainen *et al*. 2018). Therefore, we next studied whether the abundance of SAHH differs between the differentially light and temperature - acclimated kales. Separation of the proteins by SDS-PAGE and subsequent analysis by immunoblotting revealed no light-dependent adjustments in the total abundance of SAHH (Figure 9a). Analysis by native gels in turn revealed the presence of SAHH in oligomeric complexes in both Half Tall and Black Magic (Figure 9b), as previously observed in Arabidopsis leaf extracts (Rahikainen *et al*. 2017). One of the SAHH-containing complexes observed in the kale varieties corresponded to the abundant Arabidopsis SAHH complex 4 (Rahikainen *et al*., 2017), as deduced from its migration on CN-gels (Figure 9b). In Arabidopsis, increased abundance of SAHH complex 4 correlated with increased abundance of 4MO-I3M (Rahikainen *et al*. 2018). In kales, this oligomeric composition was present as a protein band doublet, comprising a prominent high MW band and a less abundant low MW band (Figure 9b). In the HL-acclimated kales, the lower MW Complex 4 band was barely detectable (Figure 9b).

**Figure 9.**
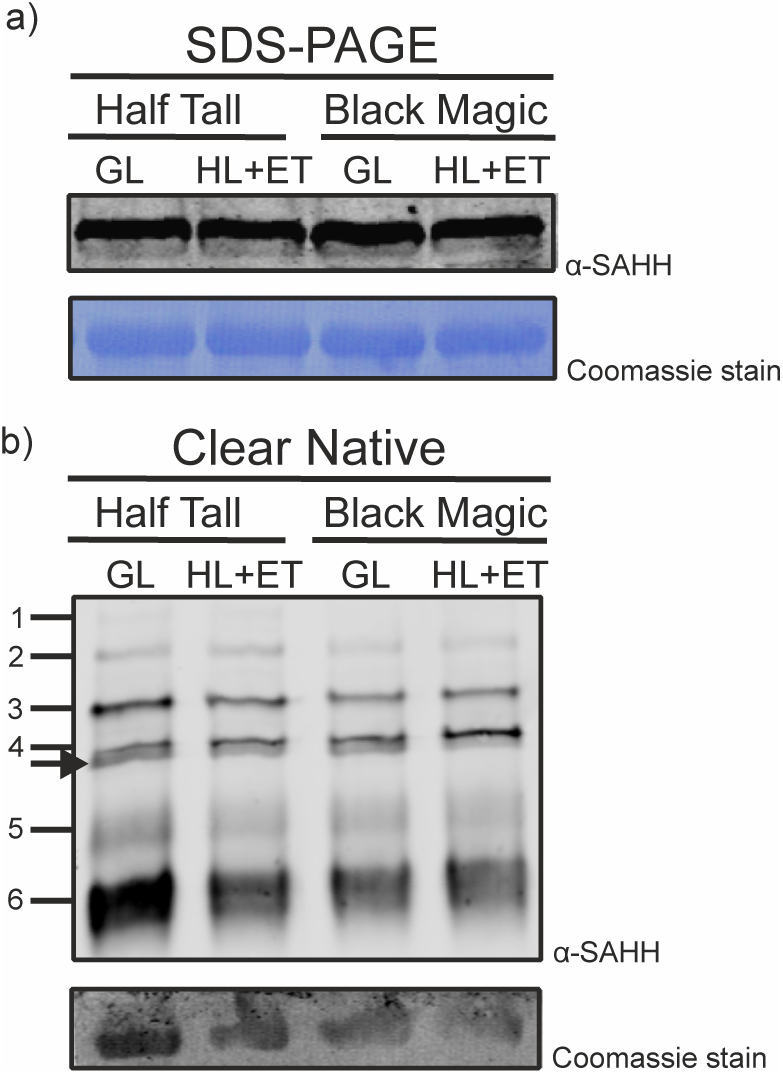
SAHH abundance and complex formation in differentially light-acclimated kale leaves. A) SAHH abundance in Half Tall and Black Magic leaves as detected by SDS-PAGE and immunoblotting with specific anti-SAHH antibody. B) SAHH-containing oligomeric complexes in Half Tall and Black Magic leaves as determined by CN-PAGE and subsequently immunoblotting with specific anti-SAHH antibody. GL, growth light; HL+ET, high light and elevated temperature.

### Glucosinolate profiles of differentially high-light-acclimated kales

Next we analysed foliar GSL profiles in differentially light-acclimated kales. The analysis detected three indole GSLs, eight aliphatic GSLs and one benzenic GSL compound. The unmodified indole GSL indolyl-3-ylmethyl GSL (I3M; glucobrassicin) can be hydroxylated in position 1 or 4 (Pfalz *et al*. 2011), forming metabolic intermediates that are subsequently methylated by indole GSL methyltransferases (IGMTs), generating the modified indole GSLs NMO-I3M (*N*-methoxy-indol-3-yl-methyl GSL) and 4MO-I3M (4-methoxy-indol-3-yl-methyl GSL) (Pfalz *et al*. 2016).

The contents of indole GSLs showed both light intensity-related and cultivar-dependent changes. Black Magic showed an overall higher content of indole GSL, which declined under long-term HL+ET irradiation (Figure 10a). On the contrary, the total content of indole GSLs in Half Tall did not display HL+ET-dependent changes (Figure 10a). Interestingly, the level of 4MO-I3M decreased upon HL+ET acclimation in both varieties (Figure 10c). The level of NMO-I3M in Black Magic showed an opposite trend with respect to 4MO-I3M, with a significant increase in HL+ET-acclimated Black Magic leaves (Figure 10d).

**Figure 10.**
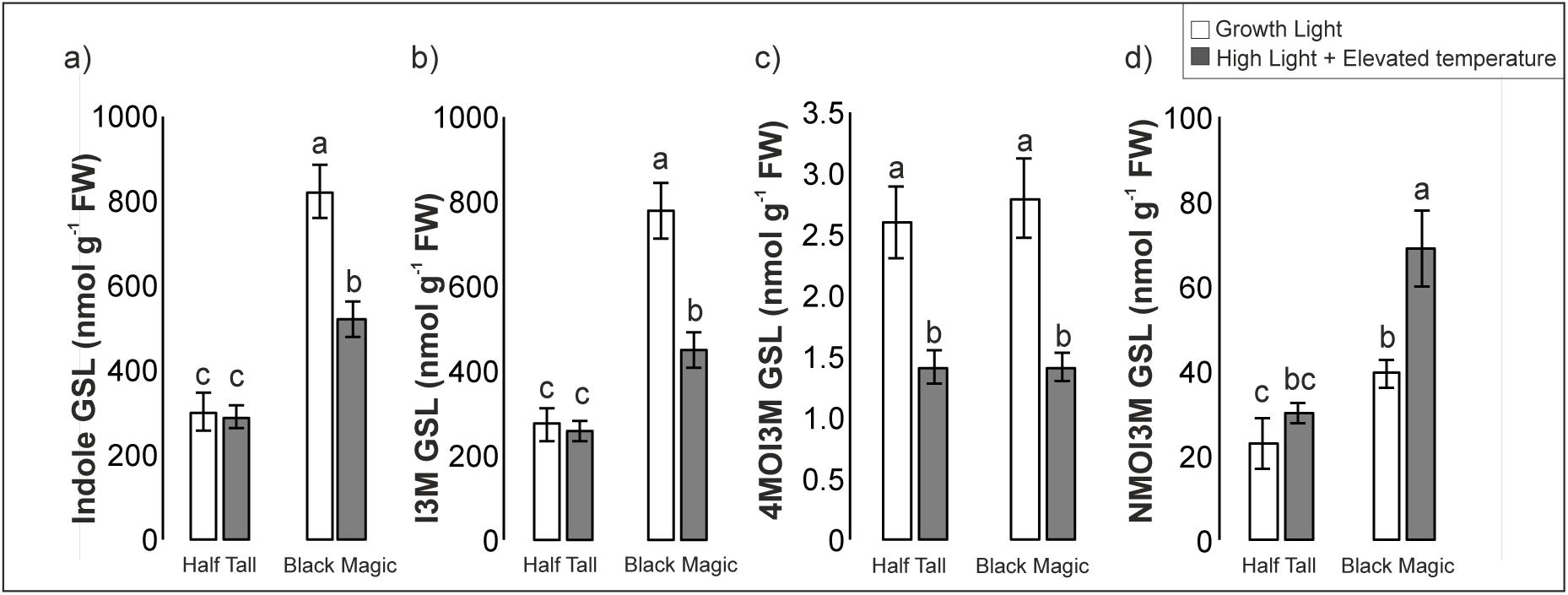
Contents of indole GSL in differentially light-acclimated kales. Half Tall (HT) and Black Magic (BM) were grown under 130 µmol photons m^−2^s^−1^ at 22°C (GL) or 800 µmol photons m^−2^s^−1^ at 26°C (HL+ET) and indole GSL were analysed by LC-MS/MS. A) Total amount of indole GSLs. B) Content of I3M (indol-3-ylmethyl GSL; glucobrassicin) C) Content of 4MO-I3M (4-methoxyindol-3-ylmethyl GSL; 4-methoxyglucobrassicin) D) Content of NMO-I3M (*N*-methoxyindol-3-ylmethyl GSL; neoglucobrassicin)

The overall content of aliphatic methionine-derived GSL increased upon HL+ET acclimation in both Half Tall and Black Magic, but the individual GSL profiles showed significant cultivar-dependent differences (Figure 11a). The biosynthesis of aliphatic GSL comprises three main steps: elongation of the amino acid chain, formation of the core GSL structure and modification of the side chain. As expected for Brassica species (Verkerk *et al*. 2009), aliphatic GSLs with C3-C5, referring to the number of carbons in their aliphatic side chains, were detected (Figure 11). Half Tall showed high contents of the C3 aliphatic GSLs 3-methylthiopropyl GSL (3MTP; glucoiberverin), 3-methylsulfinylpropyl GSL (3MSP; glucocheirolin) and 2-propenyl GSL (2PROP GSL; sinigrin), which were barely detectable in Black Magic (Figure 11c). In addition, 3-butenyl GSL (3BUT GSL) and 2(*R*)-2-hydroxy-3-butenyl GSL (2R-OH-3BUT GSL; progoitrin) were abundant in Half Tall but not in Black Magic (Figure 11d).

**Figure 11.**
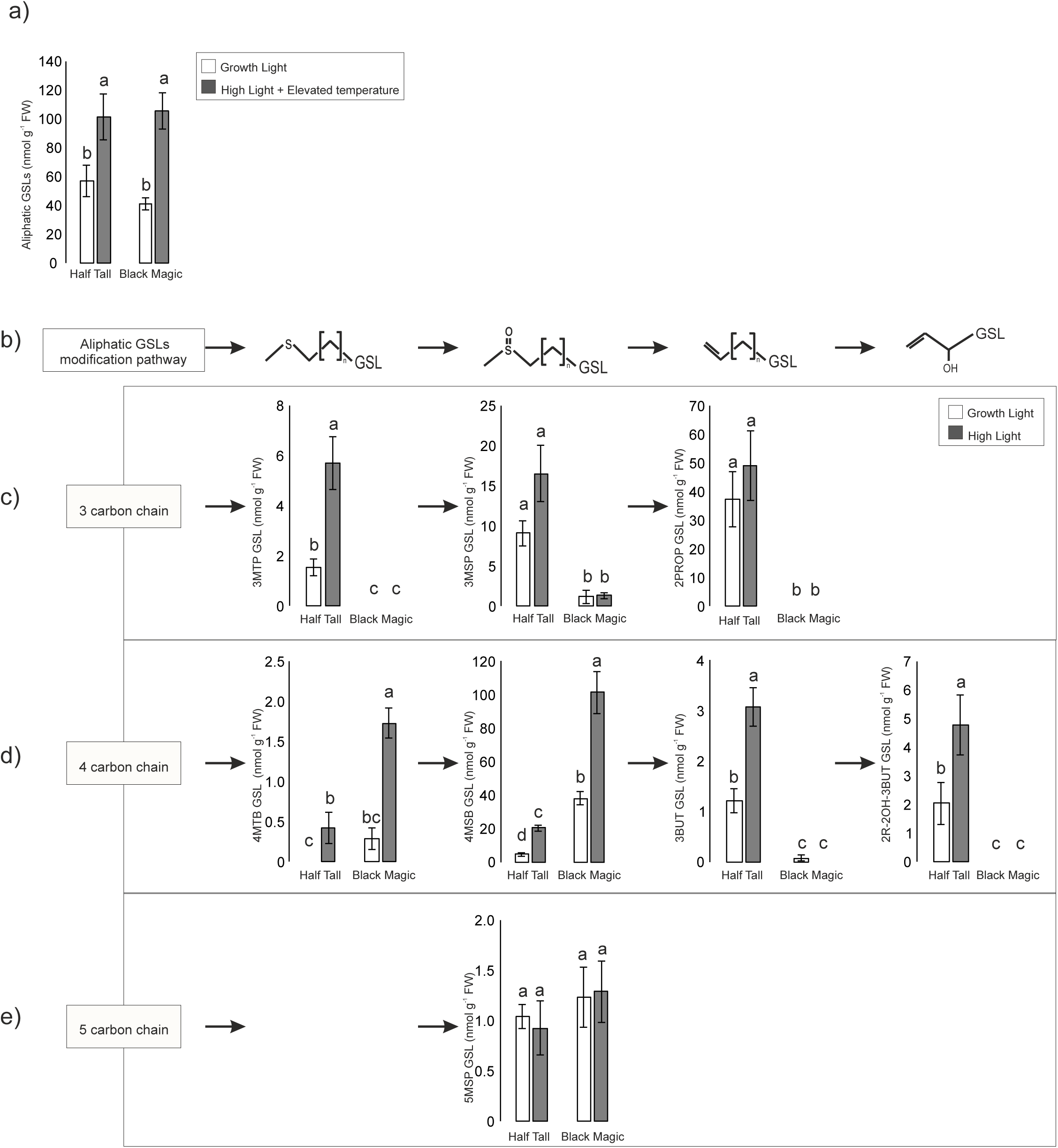
Contents of aliphatic GSL in differentially high-light-acclimated kales. Half Tall and Black Magic were grown under 130 µmol photons m^−2^s^−1^ at 22°C (GL) or 800 µmol photons m^−2^s^−1^ at 26°C (HL+ET) and aliphatic GSL were analysed by LC-MS/MS. A) Total amount of aliphatic GSL B) Schematic representation of structural modifications occurring in the GSL structure C) Content of 3-carbon side chain GSLs. 3MTP (3-methylthiopropyl GSL), 3MSP (3-methylsulfinylpropyl GSL; glucoiberin) and 2PROP GSL (2-propenyl GSL; sinigrin) D) Content of 4-carbon side chain GSLs. 4MTB (4-methylthiolbutyl GSL; glucoerucin), 4MSB (4-methylsulfinylbutyl GSL; glucoraphanin), 3BUT (2(*R*)-2-hydroxy-3-butenyl GSL; Progoitrin) E) Content of 5-carbon side chain GSLs. 5MSP (5-methylsulfinylpentyl GSL; glucoalyssin)

In contrast to Half Tall, Black Magic contained the C4 aliphatic GSL 4-methylthiobutyl GSL (4MTB GSL; glucoerucin) and was particularly rich in 4-methylsulfinylbutyl GSL (4MSB GSL; glucoraphanin) (Figure 11d). In addition, Black Magic accumulated high level of the aromatic 2-phenylethyl GSL (2PE GSL; gluconasturtiin), especially upon acclimation to HL+ET (Figure 12). The content of 5-methylthiopentyl GSL (5MSP GSL; glucoberteroin), in turn, did not show cultivar- or light intensity-dependent adjustments (Figure 11e). These findings suggest that HL+ET acclimation promotes the accumulation of aliphatic GSL, irrespective of the nature of the abundant GSL species, which is highly cultivar-dependent.

**Figure 12.**
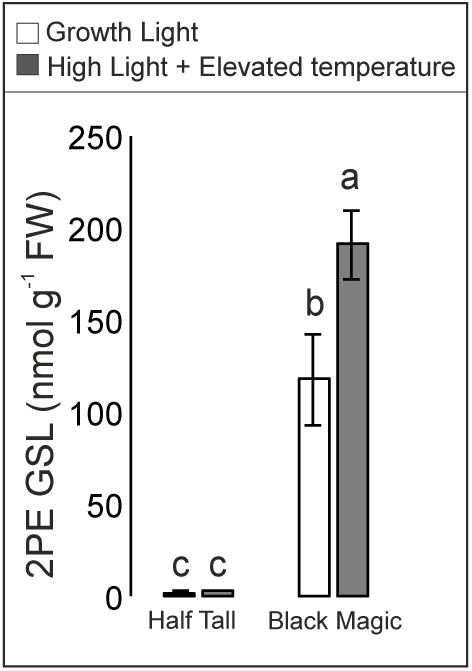
Content of the benzenic GSL 2-Phenylethyl (2PE) in differentially high-light-acclimated kales. Half Tall and Black Magic were grown under 130 µmol photons m^−2^s^−1^ at 22°C (GL) or 800 µmol photons m^−2^s^−1^ at 26°C (HL+ET) and benzenic GSLs were analysed by LC-MS/MS.

## Discussion

Light intensities that exceed the photosynthetic capacity of a plant in a given environment may result in imbalanced accumulation of redox-active intermediates and cause damage to the photosynthetic protein complexes (Aro, Virgin & Andersson 1993; Muller 2001; Miyake 2010; Kono, Noguchi & Terashima 2014; Tiwari *et al*. 2016; Gu *et al*. 2017). Photosynthetic organisms therefore undergo coordinated adjustments in gene expression and metabolism to optimize their fitness in the prevailing growth environment. Under natural conditions, fluctuations in light intensity are commonly seen as an environmental stress factor, while in greenhouse conditions, artificial manipulation of plant metabolomes by alterations in growth conditions may allow improved production of desired end products. However, the basic understanding of how the growth conditions affect the chemical composition of crops is still limited. In this study, we aimed towards understanding how long-term growth under a combination of high irradiance and elevated temperature affects the chemical composition and growth in differentially pigmented varieties of kale.

### Long-term plant acclimation to high light involves reprogramming of gene expression and protective metabolism

Light-induced acclimation responses depend on the severity and the duration of excess irradiation. A number of studies have elaborated the transient nature of HL-induced changes in foliar transcriptomes (Vogel *et al*. 2014; Crisp *et al*. 2017), while long-term effects on transcriptomic adjustments have drawn less attention. Here we assessed the transcript profile of HL+ET-acclimated Arabidopsis leaves (Supplementary Table S1), and compared the acclimation response to previously reported short-term HL-induced transcriptional adjustments (Kleine, Kindgren, Benedict, Hendrickson & Strand 2007; Jung *et al*. 2013; Gläßer *et al*. 2014; Schmitz *et al*. 2014) to elucidate how long-term growth under HL affects physiological processes in Arabidopsis.

Long-term HL+ET acclimation of Arabidopsis was accompanied by increased abundance of transcripts related to the biosynthesis of flavonoids and anthocyanins, and reduced transcript abundance for proteins involved in photosynthetic light harvesting, when compared to plants grown under moderate GL (Figure 1b). This protective response was evident also as visually observable accumulation of blue and purple pigments, which accumulate in leaf epidermal cells to protect HL+ET-acclimated leaves against light stress (Figure 1a; Chalker-Scott, 1999). A distinguishing feature between long-term HL+ET-acclimated leaves and short-term light stressed ones was reduced abundance of transcripts in GO categories related to biotic stress responses, in comparison to non-stressful GL conditions (Table S11), suggesting alleviation of defense priming and suppression of hypersensitive cell death in leaves acclimated to the potentially stressful abiotic cues. Long-term morphological adjustments, including development of thick leaves rich in anthocyanins and other phenolic metabolites, may be deterrent against pathogens and herbivores and promote cross-tolerance against biotic stress agents.

Higher levels of carotenoids and anthocyanins were detected in Black Magic kale, in comparison to Half Tall, and these were shown to increase significantly after growth under HL+ET (Figure 6). Upregulation of the protective pigments correlated with increased photosynthetic electron transport under high irradiance in Black Magic, which was also improved after HL+ET growth, which can be attributed to lower levels of NPQ in Black Magic (Figure 5). Since many of the protective metabolites have health-promoting nutritional effects in humans (Verkerk *et al*. 2009; Dinkova-Kostova & Kostov 2012), light-induced adjustments in foliar chemical composition can directly impact the nutritional value of leafy vegetables. Therefore, long-term exposure to warm high light conditions can be applied as a tool to trigger the production of health- and taste-related compounds with the aim to increase the commercial value of crops in greenhouse cultivation.

### The kale varieties contain increased amounts of genotype-dependent aliphatic glucosinolate structures when grown under high light and elevated temperature

GSLs are major defensive compounds commonly associated with plant-biotic interactions in the order Brassicales (Halkier & Gershenzon 2006; Hopkins, van Dam & van Loon 2009). To date, around 130 GSL species have been identified and their occurrence in various plant cells, tissues (Agerbirk & Olsen 2012) and species under different developmental stages (Brown, Tokuhisa, Reichelt & Gershenzon 2003) and environmental conditions (Cartea, Velasco, Obregón, Padilla & de Haro 2008; Huseby *et al*. 2013; Martínez-Ballesta, Moreno & Carvajal 2013) has been extensively characterized. Metabolite profiling of the Half Tall and Black Magic varieties of kale revealed that ambient growth light and temperature can significantly affect the contents of both indole and aliphatic GSLs in kale (Figures 10 and 11).

The methionine-derived aliphatic GSLs form a predominant group of natural compounds in *A. thaliana* and many crops in the Brassicaceae family. We show that the Half Tall and Black Magic varieties of kale undergo a HL-induced increase in total aliphatic GSL levels, although the individual GSL species display cultivar-dependent changes (Figure 11). Black Magic accumulated aliphatic GSLs with a length of 4 carbons in their aliphatic side chain, while aliphatic side chains of 3 carbons predominated in Half Tall, presumably due to occurrence of different biosynthetic machineries in these two varieties.

The committed enzyme responsible for the chain elongation step in the biosynthesis of aliphatic GSLs is methylthioalkylmalate synthase (MAM; Kliebenstein *et al*., 2001; Kroymann, 2001). The length of the aliphatic side-chain is determined by the number of times it undergoes a MAM-driven elongation (Kliebenstein *et al*. 2001b; Kroymann 2001), but it is notable that MAM isoforms elongate the methionine-derived keto-acids in a chain-length-dependent manner. The profile of aliphatic GSL species with different chain length is therefore essentially determined by the MAM isoforms that are present in a given plant species, cultivar or ecotype (Kroymann, Donnerhacke, Schnabelrauch & Mitchell-Olds 2003; Kroymann *et al*. 2006). In Arabidopsis, MAM is present as three different isoforms, MAM1, MAM2 and MAM3 (Benderoth, Pfalz & Kroymann 2009). MAM2 is responsible for the first elongation cycle of methionine and forms 3-carbon aliphatic side-chain, whereas MAM1 is able to generate both 3-carbon and 4-carbon aliphatic side-chains. The differential profiles of aliphatic GSLs in Half Tall and Black Magic (Figure 11b,c) suggest that the kale varieties possess different isoforms of MAM. The lack of 3-carbon aliphatic side-chains in Black Magic suggests that this variety is devoid of MAM2 (Figure 11c,d). In contrast, in Half Tall that accumulates 3-carbon aliphatic side-chain GSL, an enzyme with the biochemical properties of the Arabidopsis MAM2 must be present. Contrastingly, high content of the 4-carbon aliphatic side-chain GSL 4MSP in Black Magic (Figure 11c,d) suggests the occurrence of a MAM1-like enzyme. Altogether, while the growth light intensity seemingly modulates the overall accumulation of GSLs, the side-chain carbon length of the aliphatic GSLs is genotype dependent.

Another relevant polymorphic locus controlling GSL profiles is *GSL-AOP* (Magrath *et al*. 1994; Mithen, Clarke, Lister & Dean 1995). It operates downstream of biosynthesis of the GSL core structure, and its presence or absence determines whether a given species or cultivar predominantly accumulates hydroxyalkyl GSLs, alkenyl GSLs or methylsulfinyl GSLs (Kliebenstein *et al*., 2001a,b, 2007). In Half Tall, the presence of 2-propenyl GSL (alkenyl GSL) pointed to the presence of a functional AOP2 in this variety (Figure 8c). In contrast, Black Magic acumulated methylsulfinyl GSL in the form of 4-methylsulfinylbutyl GSL (Figure 8d), which is not further converted to other glucosinolate structures, due to absence of the AOP enzymes in this variety.

The biosynthetic machineries behind aliphatic GSL biosynthesis are drawing increasing research interest, as understanding the molecular machineries responsible for the enormous GSL diversity may pave the way for traditional breeding or biotechnological manipulation of GSL content and their pungent metabolites in Brassica crops (Petersen, Wang, Crocoll & Halkier 2018; Kumar *et al*. 2019). In this study, we provide evidence indicating that besides the evolutionary and biochemical foundations of GSL metabolism (Kumar *et al*. 2019), optimized light conditions can be applied to modulate the GSL profiles to increase the contents of beneficial GSL compounds while decreasing those with deleterious effects.

### High light stress as a noninvasive means for cultivation of healthier plants

The biosynthetic machineries of plants are highly responsive to light and their metabolite profiles can therefore be non-invasively manipulated by changes in the intensity and spectral quality of light (Cargnel, Demkura & Ballaré 2014). In greenhouse cultivation, manipulation of metabolic and developmental processes by alterations in growth conditions can therefore be applied to enhance the production of desired compounds. GSLs have beneficial nutritional effects but detrimental effects have also been reported (Greer & Deeney 1959; Tripathi & Mishra 2007; Romeo, Iori, Rollin, Bramanti & Mazzon 2018; Tafakh *et al*. 2019).

In this study, GSL profiling provided insights to differential nutritional qualities of kale varieties. The purple variety, Black Magic, was rich in the health-promoting 4MSB GSL (glucoraphinin) and 2PE (glucocasturtiin), and their contents became further increased when the plants were grown under HL (Figures 11d and 12). Among GSL structures with potential harmful effects, progoitrin, also known as 2*R*-2-OH-3-butenyl GSL, has been associated with bitter taste and long intake periods may cause goiter in animals (Greer 1957; Felker *et al*. 2016). It is therefore notable, that in Black Magic, the levels of 2*R*-2-OH-3-butenyl GSL were below detection in both light conditions studied (Figure 11d). In contrast, Half Tall responded to HL growth by accumulating 2*R*-2-OH-3-butenyl GSL (Figure 11d).

Several GSL structures and their breakdown products have been reported beneficial in human diet (Traka & Mithen 2009; Gupta *et al*. 2015; Lee *et al*. 2019). Among indole GSLs, degradation of 4MO-I3M yields indole-3-carbinol (I3C), which was recently demonstrated to have anti-cancerous activity through inhibition of the HECT-type E3 ubiquitin ligase WWP1 that promotes tumorigenesis in several cell types (Lee *et al*. 2019). Enzymatic processing of the aliphatic 4MSB GSL and 2PE GSL into sulphoraphane and phenethyl isothiocyanate (PEITC), respectively, yields metabolites with health-beneficial properties through anticarcinogenic and chemo-protectant activities (Cheung & Kong 2010; Jiang *et al*. 2018). These GSL-derived metabolites can reduce the amounts of carcinogens by inhibiting phase I enzymes and activating phase II enzymes (Eastham, Howard, Balachandran, Pasco & Claudio 2018). Moreover, growth-inhibitory effects for sulforaphane in head, neck and prostate cancer cells have been reported (Singh *et al*. 2005; Gupta *et al*. 2015) and PEITC has been associated with prevention of the growth of oral cancer cell lines (Chen *et al*. 2012). Currently, their potential effects on different cancer types is a matter of extensive exploration (Castro *et al*. 2019; Mitsiogianni *et al*. 2019; Upadhyaya, Liu & Dey 2019; Yin *et al*. 2019). Sulphoraphane has also been proposed as a potential therapy for precluding vascular complications in diabetes (Yamagishi & Matsui 2016). Beneforté® Broccoli, highly enriched in 4MSB-GSL, is commercially available worldwide and crop industry continues prompting research towards generation of bio-refined crops.

Growth of Black Magic under high irradiance promoted the accumulation of health-promoting aliphatic GSLs and anthocyanins, while the disadvantageous GSL structures remained below detection limits (Figures 6 and 11). On the other hand, reflecting the reduced transcript abundance for genes related to methoxylation of indole GSLs in the Arabidopsis long-term HL transcriptome, both Black Magic and Half Tall displayed reduced levels of the indolic 4MO-I3M when grown under HL+ET (Figure 10). Altogether, growth light intensity is a key factor that can impact the accumulation of beneficial metabolites in commercially valuable varieties of Brassica species.

**Table 1: Anthocyanins content in Black Magic.**

No.: number of the peak in the obtained chromatograms (depicted in Figure S3); RT_min_: retention time in minutes; [M+ H]^+^: monoisotopic mass; Tentative ID: tentative identification.

## Acknowledgements

This work was supported by Academy of Finland project 307719 to S.K., 325122 to JP, 303757 to E-M.A. and the Academy of Finland Center of Excellence in Primary Producers 2014-2019 (307335). BY and WY acknowledge the funding for Competitive funding to strengthen universities’ research profiles (Profi 4, Decision No. 318894, Subproject: FOOD INNOVATIONS) by the Academy of Finland. S.A. was funded by the University of Turku Doctoral Programme in Molecular Life Sciences. MB was funded by the Danish National Research Foundation, DNRF (grant 99). The authors declare no conflict of interest.

