## Supplemental Figures for "Growth under high light and elevated temperature affects metabolic responses and accumulation of health-promoting metabolites in kale varieties"

**Supplementary data legends.**

**Supplementary Figure 1. GO enrichment categories trees from the 7 clusters derived from the hierarchical clustering analysis.** The 30 most significant GO categories were used.

A) GO enrichment tree from cluster 1.

B) GO enrichment tree from cluster 2.

C) GO enrichment tree from cluster 3.

D) GO enrichment tree from cluster 5.

E) GO enrichment tree from cluster 6.

F) GO enrichment tree from cluster 7.

**Supplementary Figure 2. GO enrichment categories in the Long-term high light comparison analysis tree.** The 30 most significant GO categories were used.

A) Up-regulated GO enrichment tree

B) Down-regulated GO enrichment tree.

**Supplementary Figure 3. Representative chromatogram of the LC-MS analysis of Black Magic anthocyanins.** Orders refer to retention time, ordinates refer to absorption intensity. Peaks position are marked with number 1 to 10, and their tentative identities are listed in Table 1.

### Supplemental Figure 1

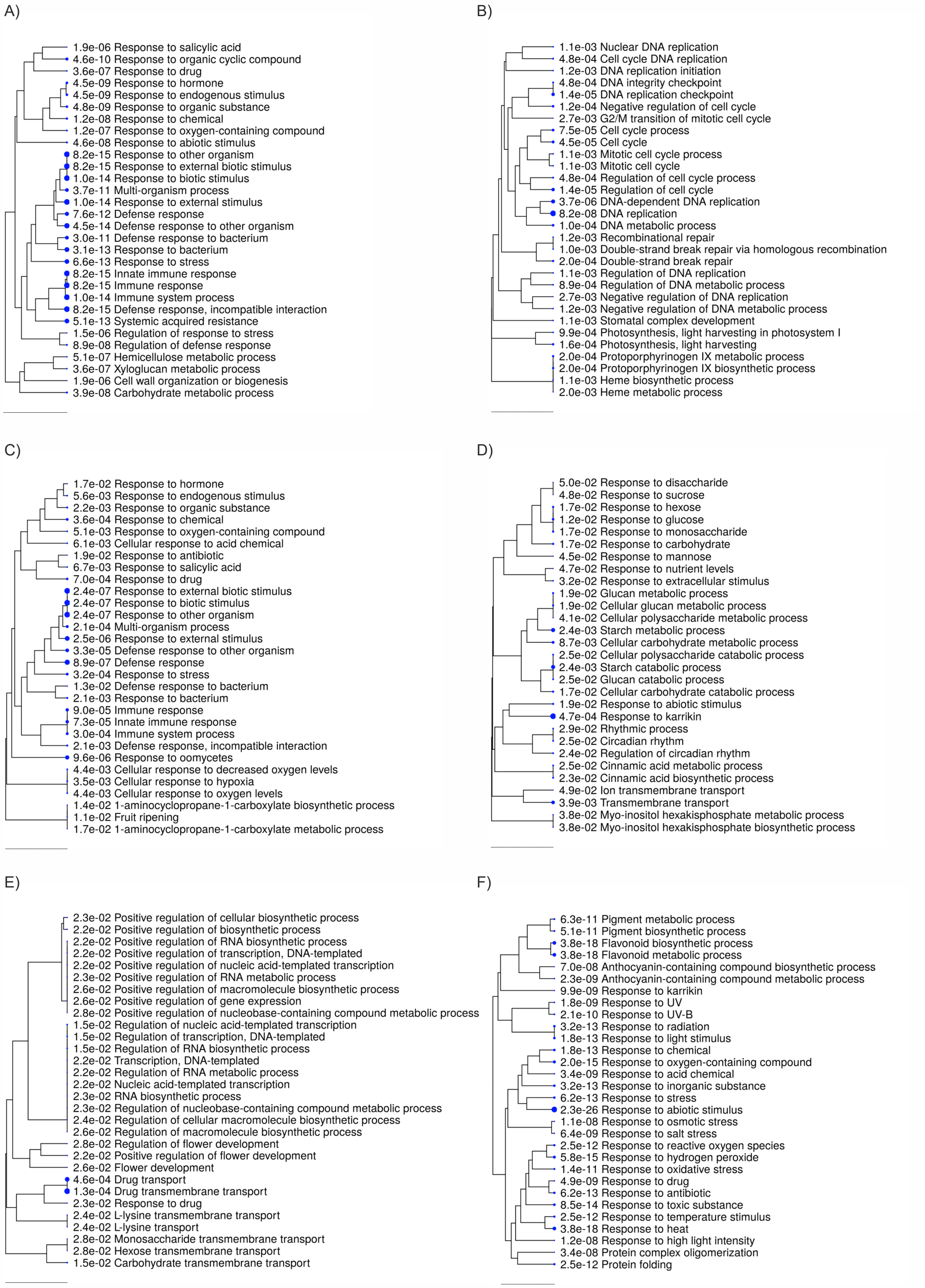

Supplemental Figure 2

A)

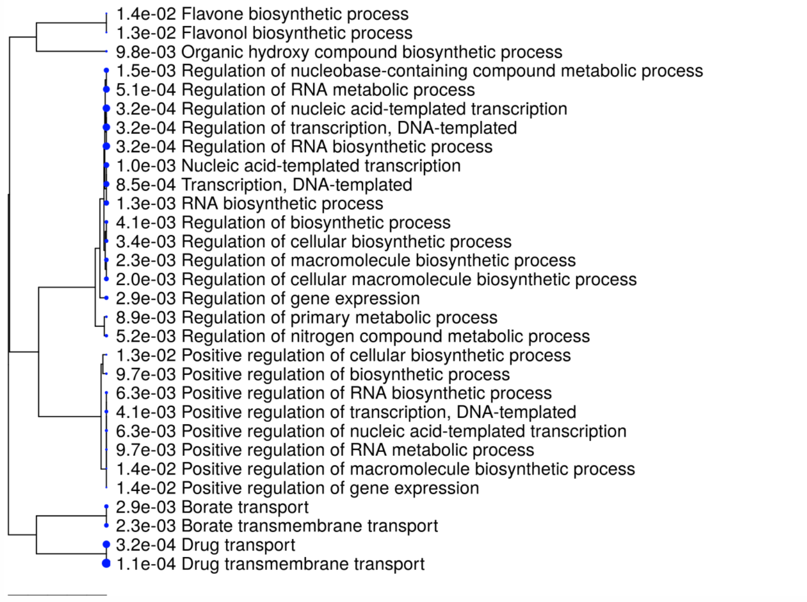


B)

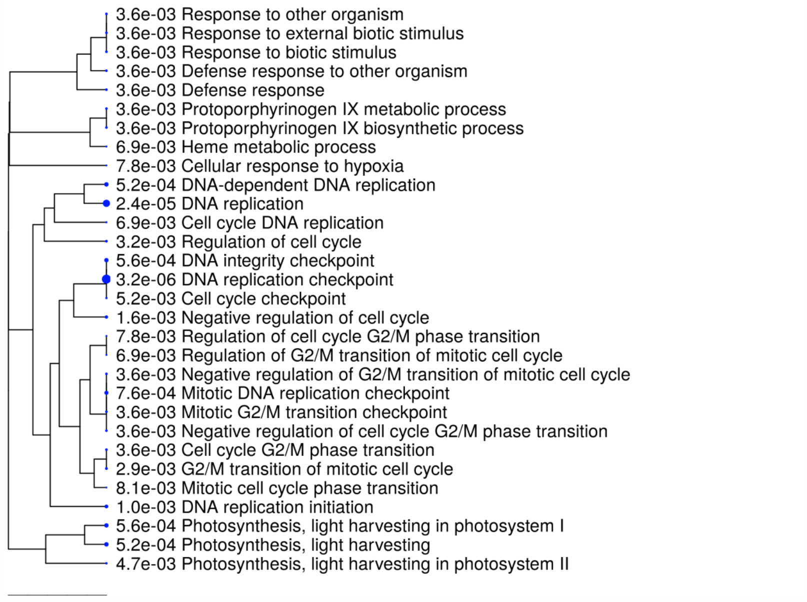

Supplemental Figure 3

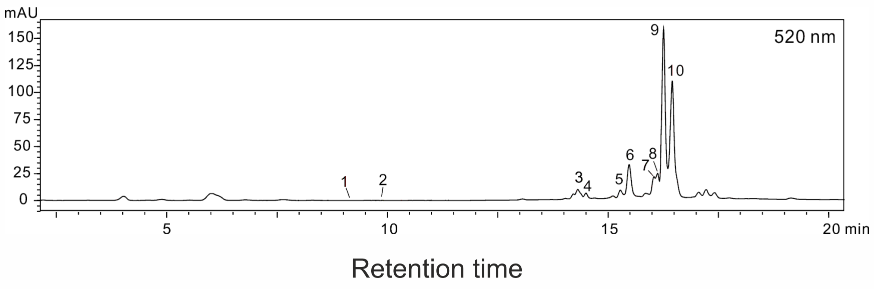
